# One-Step PCR Amplicon Library Construction (OSPALC, version 1)

**DOI:** 10.1101/2022.10.26.511284

**Authors:** Jiahao Ni, Jiao Pan, Yaohai Wang, Tianhao Chen, Xinshi Feng, Yichen Li, Tongtong Lin, Michael Lynch, Hongan Long, Weiyi Li

## Abstract

High-throughput sequencing of amplicons has been widely used to precisely and efficiently identify species compositions and analyze community structures, greatly promoting biological studies involving large amounts of complex samples, especially those involving environmental and pathogen-monitoring ones. However, commercial library preparation kits for amplicon sequencing, which generally require multiple steps, including adapter ligation and indexing, are expensive and time-consuming, especially for applications at a large scale. Here, to overcome this technical hurdle, we present a one-step PCR amplicon library construction (OSPALC) protocol for amplicon library preparations in the lab along with a QIIME2-based amplicon analysis pipeline. High-quality reads have been generated by this approach to reliably identify species compositions of mock bacterial communities and environmental samples. With this protocol, the amplicon library construction is completed through one regular PCR with long primers, and the total cost per DNA/cDNA sample decreases to just 1/15 of the typical cost via service providers. Empirically tested primers to construct OSPALC libraries for 16S rDNA V4 regions are demonstrated as a case study. Criteria to design primers targeting at any regions of prokaryotic and eukaryotic genomes are also suggested. In principle, OSPALC can be conveniently used in amplicon library constructions of any target gene of both prokaryotes and eukaryotes at the DNA or RNA levels, and will facilitate research in numerous fields.

## Introduction

Amplicon sequencing of single gene fragments is widely used for studying biodiversity and community compositions, based on reference databases and relative abundance of the target genes. It can be conveniently applied to track tempo-spatial variations of population and community structures, resulting from genetic and/or environmental changes (Beser et al. 2017; Costello et al. 2009; Gous et al. 2019; McNamara et al. 2020; Pochon et al. 2013; Song et al. 2019; Xu et al. 2012). Accumulated amplicon sequences of various samples also provide growing databases for future research. With the development of sequencing technology and advanced analytical tools, amplicon sequencing has been adapted to various sequencing platforms, from short-reads technology such as Illumina, Roche 454, Ion Torrent to Nanopore/PacBio long-read sequencing (Beser et al. 2017; Bokulich et al. 2018; Bolyen et al. 2019; Costello et al. 2009; Fadrosh et al. 2014; Moonsamy et al. 2013; Myer et al. 2016; Neiman et al. 2011; Robeson et al. 2020; Wu et al. 2015). Although widely applicable, costs for amplicon library preparations of a large number of samples are much higher than those for the downstream sequencing. Even higher costs are incurred if library constructions and data analyses are outsourced to a service provider.

Illumina paired-end sequencing (PE150 and PE250) has become the most widely-used amplicon sequencing technology. A series of library preparation kits are available, such as QIAseq 1Step Amplicon Lib UDI-A Kit (Cat. No.: 180419), Vazyme VAHTS AmpSeq Library Prep Kit V2 (Cat. No.: NA201), and so on. Protocols for these kits usually contain multiple procedures such as the adapter ligation and a further PCR for indexing (Neiman et al. 2011; Vo and Jedlicka 2014). Such protocols are always time-consuming and costly, due to the complex procedures, which require expensive reagents. In addition, the two contiguous PCRs can lead to severe problems, such as primer dimers, sample contaminations and biased amplifications (Bohmann et al. 2022). Although promising, recently-proposed full-length amplicon sequencing by PacBio or Nanopore long-read platforms also suffers from these and other problems such as low reads accuracy, as well as the immature analytical tools and higher costs (Calus et al. 2018; Ciuffreda et al. 2021; Karst et al. 2018; Martijn et al. 2019; Tedersoo et al. 2021; Wagner et al. 2016).

To tackle these difficulties, we developed a fusion primer method, i.e. One-Step PCR Amplicon Library Construction (OSPALC, also a letter combination which could be unscrambled into 90 words in word games), which integrates: the design and optimization of long primers containing sequencing adaptors and primers of the target region, library construction procedures, and sequencing strategies. The method was evaluated by sequencing and analyzing 16S rDNA V4 regions of two mock communities of bacteria. Compositions of mock communities were reliably revealed by OSPALC. In addition, we were also able to apply the method, using cDNA of time-series samples, to accurately monitor the dynamics of bacterial community compositions in a summer pool, which experienced strong disturbance from hose-cleaning, as well as those of another undisturbed pond.

## Results

### Primer design and the protocol for One-Step PCR Amplicon Library Construction (OSPALC)

Each long primer for the OSPALC method is about 90 nt and contains four parts, in the 5′ to 3′ direction: the P5 or P7 sequence for Illumina flow-cell binding, 8 nt index, Illumina sequencing barcode, and the forward or reverse primer targeting at the gene of interest (Fig. 1). To minimize the chance of forming primer dimers and maximize the primer specificity, we optimized each pair of the 8-nt dual-indices by eliminating palindromes or high sequence similarity. The indices we used in this study and additional candidates, as well as criteria for selecting indices are listed in Supplemental Tables S1 and S2. The total cost for each amplicon sample starting with environmental DNA is ~$2 ($4 if starting with total RNA), after taking long-primer synthesis, PCR for library construction, and Illumina PE250 sequencing into account. These are extremely low compared with the charges by service providers, for example, ~$30 or most costs for DNA samples. Many service providers do not even accept cDNA or RNA samples.

**Figure 1.**
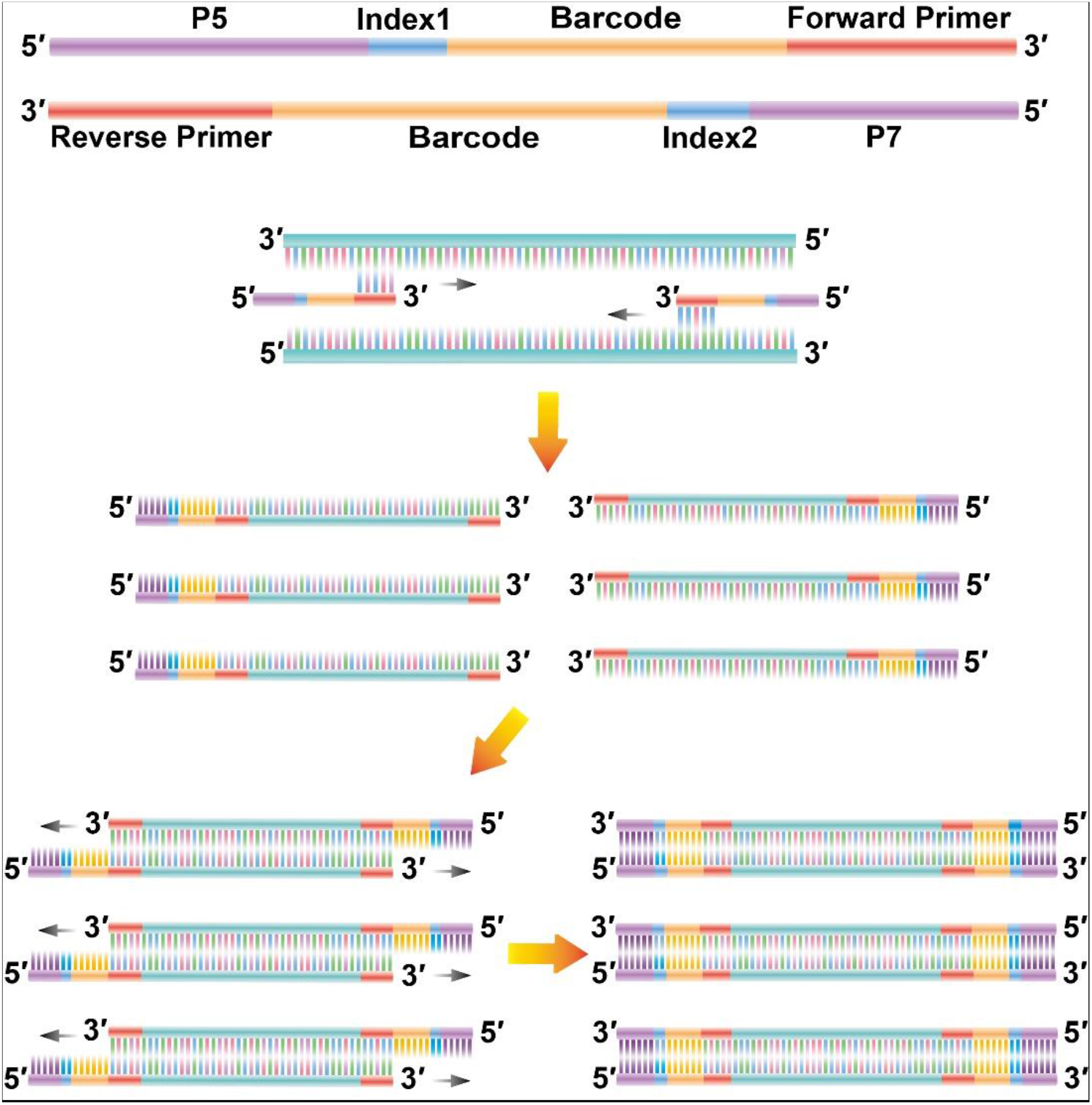
Long primers and PCR for OSPALC. The top two horizontal stacked bars show the structures of long primers. Each long primer contains four parts: P5/P7, for the flow cell binding; index1/index2, for sample-indexing; barcode, for sequencing enzyme binding; forward/reverse primer, for amplification of the target gene region. The steps below the long primers illustrate the PCR procedures for OSPLAC.

The OSPALC procedures (literally, just one regular PCR with long primers; after testing different PCR conditions as shown in the next section; Supplemental Table S3) are as follows:

Any high-fidelity PCR kit could be applied, and here we used 2× Phanta Flash Master Mix (Dye Plus) (Vazyme Inc., Cat. No.: P520-02). The 50 μl reaction system consisted of 1 μl DNA template (10 ng/μl), 25 μl master mix, 1 μl of the forward long primer (10 μM), 1 μl of the reverse long primer (10 μM) and 22 μl sterilized ultra-pure water, with 10 regular and 10 gradient PCR cycles (a miniaturized 10-μl protocol is provided in Supplemental File S1). Unless the concentration of the pooled-libraries is lower than the requirement for the sequencer, as few PCR cycles as possible are recommended. One blank control with sterilized DI water as DNA template is also preferred.

Each library—amplified gene fragments with adaptors on—of 400 to 500 bp, was then size-selected by gel cutting and purified with the E.Z.N.A.^®^ Gel Extraction Kit (OMEGA, Cat. No.: D2500-02; preferred over the size selection with magnetic beads). If the size distribution of amplicon libraries looks uniform on the agarose gel, we recommend to perform the size selection on the pooled libraries, which could significantly reduce the workload in the gel extraction step. Illumina PE250 sequencing on the pooled amplicon libraries was then performed. PE150 could also be applied, if the size of the target gene fragment is short enough.

### Compositions of the two mock communities are reliably revealed by OSPALC vs. methods by service providers

To evaluate the data quality from OSPALC, we first prepared two mock communities by mixing purified genomic DNA of five bacterial species, including three Gram-negative species: *Escherichia coli* MG1655, *Pseudomonas aeruginosa* PAO1, *Shewanella putrefaciens* CGMCC1.6515, and two Gram-positive ones: *Bacillus subtilis* ATCC6051 and *Kocuria polaris* CGMCC1.8013. One mock community (Mock Equal—ME) contained genomic DNA with equal copy numbers of 16S rDNA in each species, by taking account of the specific 16S rDNA copy number of each bacterial genome. The other (Mock Gradient—MG) contained genomic DNA with gradients of 16S rDNA copy number of the five bacteria (Supplemental Table S4). We then constructed amplicon libraries from the 16S rDNA V4 region for the mock communities using OSPALC. As a comparison, we outsourced the genomic DNA samples of the same mock communities to the service provider in two batches, where amplicon libraries were constructed with kits including end repair, adapter ligation and indexing in three separate PCR steps, as well as multiple beads-cleaning in between (TianGen Corp.; Cat. No.: NG102). These complex steps are highly likely to increase the risk of contamination. All PE250 data were analyzed with a pipeline based on QIIME2 (version: 2021.8). After the data were imported into QIIME2, the paired-end reads were joined, and chimeric feature sequences were identified and filtered out by vsearch (Rognes et al. 2016). Joined reads were then filtered by q-score and denoised by deblur (Amir et al. 2017). Through the phylogenetic inference and the classified feature sequences, we finally obtained the species-abundance tables.

In both batches of data, analytical results of the two mock communities revealed by both OSPALC and the method of the service provider detected the five bacterial species (Fig. 2; Supplemental Fig. S1). Surprisingly, there was widespread contamination of other bacteria not belonging to the five species in the first batch of the amplicon libraries constructed by the service provider (Supplemental Fig. S1). This observation demonstrates that more library preparation steps or people touching the samples by outsourcing the amplicon library construction increase the chance of sample contamination. Also, chimeras generated during at least two PCRs for adaptor ligation, indexing or sequencing, using classical amplicon library preparation methods can be identified as other species, and thus become ‘contaminating bacteria’. These reflect another advantage of the in-lab OSPALC, since only one PCR is needed.

**Figure 2.**
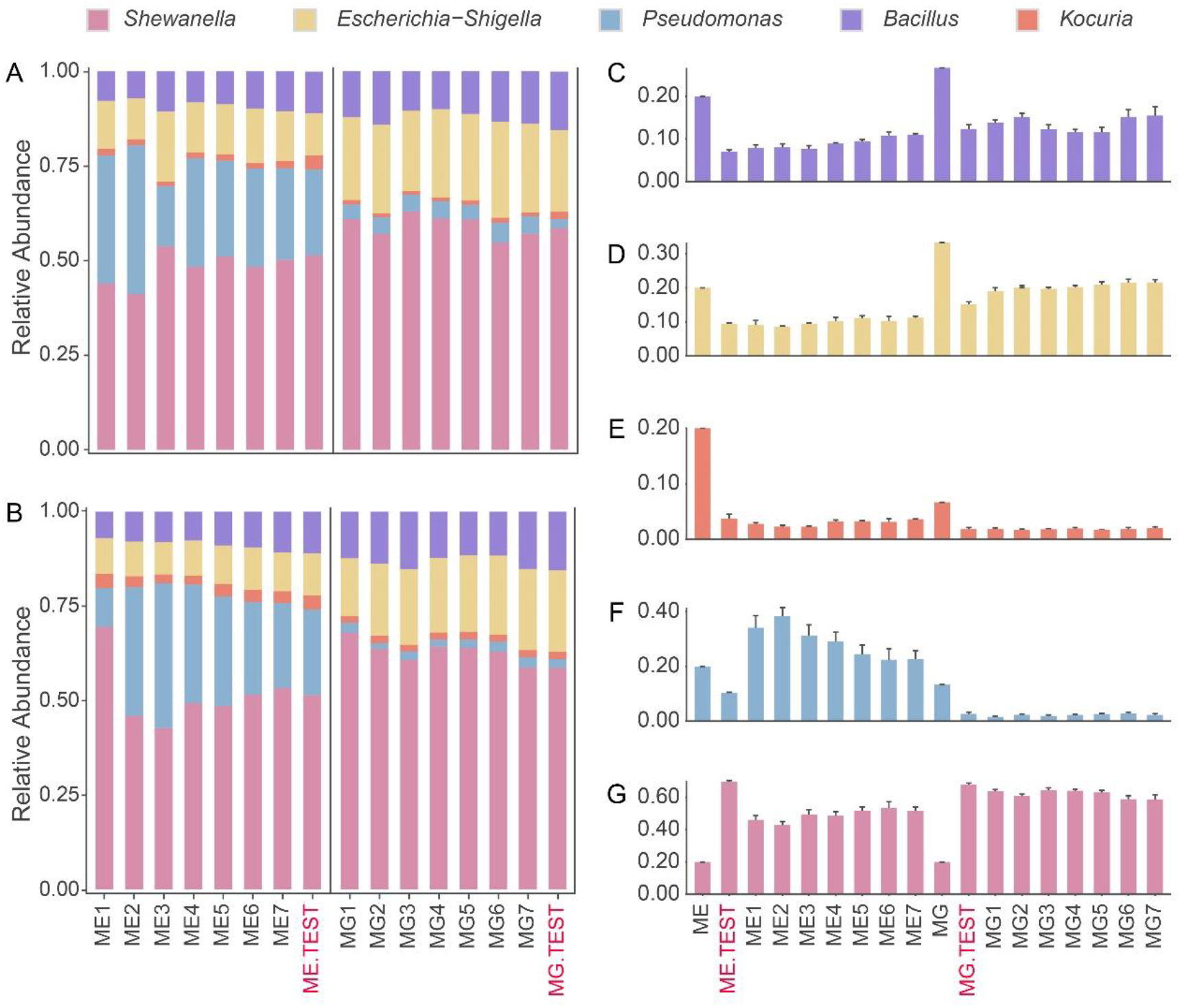
Species abundance of the mock communities, based on libraries constructed by OSPALC and the service provider. ME (Mock equal, genomic DNA with equal copy numbers of 16S rDNA in each species, the specific 16S rDNA copy number of each bacterial genome was taken into account) and MG (Mock gradient, containing genomic DNA with gradients of 16S rDNA copy number of the five bacteria, details are in Supplemental Table S4) are the known preset proportions of each bacterium in the equal and the gradient mock communities. ME1-7 and MG1-7 are the equal community and the gradient community with different PCR conditions of OSPALC. Data from the service provider are labelled with ME.TEST and MG.TEST in red. A, B: mock community sequenced with PE250 and PE150, respectively. C-G: composition variation of the five bacteria detected by PE250; the standard deviations between replicates are low and both OSPALC and service provider community compositions deviated from the preset values.

Sequencing details of the amplicon libraries by in-lab OSPALC and the service provider are in Supplemental Tables S5–S7. The service provider charged ~$30 for each amplicon sample, including the library construction and the PE250 sequencing. In comparison, the total cost using the OSPALC method, from genomic DNA to Illumina sequences is ~$2 per sample. After proceeding with the QIIME2 pipeline and PE250 reads, an average of 157,692 final joined reads per sample (8 million raw reads in total for 56 samples; joined reads passed filters against low-quality and un-joined reads) were obtained. In most cases, 50,000 joined reads per sample are sufficient for downstream analysis (sequencing fee ~$1.39 per sample). Although PE150 sequencing costs much less than PE250, we recommend PE250 sequencing. Because there is a risk that short overlaps between the PE150 forward and reverse reads result in extremely low success rate of the reads-joining step in the pipeline, especially if the insert size is close to 300 bp. In order to validate this, we also use the in-lab OSPALC libraries of the mock community samples, which are Illumina-PE150-sequenced. Sequencing the 56 samples with a total of 53.94 million raw reads, after filtering, yielded an average of 49,946 joined reads per sample, with the sequencing cost of about $0.84 per sample. However, compared with the PE250 mode, shorter read length of the PE150 mode greatly decreased the downstream join rate, i.e. only about 5% OSPALC raw reads could be joined and pass the cutoff line of 291 bp for the joined PE150 reads. Therefore, PE150 turns out not to be better, compared with the join rate of 91.78% of the PE250 reads (Supplemental Tables S5–S7). Nonetheless, the community structures revealed by both PE150 and PE250 sequencing modes are highly similar, i.e. PE150 is still useful even if only tiny part of the reads are analyzable (Fig. 2A, B). For the 16S rDNA V4 target region (~264 bp) in this study, considering the cost efficiency and potential variations in the length of amplicons, PE250 is definitely a better choice. We thus highly recommend PE250 sequencing for OSPALC libraries, especially when the target amplification region is close to or longer than 300 bp.

Species compositions of the mock communities, revealed by both the OSPALC and the service provider’s methods, show discrepancies from the preset ones because of the bias of amplification. Such bias still exists after we performed qPCR on the 16S rDNA V4 region, using genomic DNA of each bacterium for the mock community and taking account of the rDNA copy number in the genome of each bacterium (Supplemental Table S4). Nonetheless, the biases between replicates of a mock population do not significantly differ (Fig. 2; Supplemental Tables S8– S12). Such bias is a widely-known limitation for almost all amplicon methods (Bohmann et al. 2022; Yeh et al. 2021). It could be caused by multiple factors of different experimental steps, mainly the bias in the initial rounds of PCRs. To test whether the results of OSPALC could be optimized by changing the PCR conditions, we performed multiple PCRs with different reaction conditions, using both the ME and the MG mock community DNA as templates, each with four replicates (Supplemental Table S3). In detail, seven sets of PCR conditions were used to construct OSPALC libraries. In the four PCR sets, we set up the annealing temperature of 55 °C with 35, 30, 25 and 20 amplification cycles (labelled as 1 to 4, respectively). In the other three treatments (5 to 7), the annealing temperature was gradually increased from 55 °C to 65 °C after the tenth cycle with a total of 20, 25 and 30 cycles. This gradient of annealing temperature was set up to increase the primer-binding specificity because we expect the whole long primers to fully pair with the templates in later rounds of amplifications. The results from all the above trials did not show any significant differences (Fig. 2A, B), the CV (coefficient of variation) of the deviations from the genuine community compositions among replicates is quite low, indicating that the discrepancy from the real community compositions may result from the amplification bias generated from the original 10 PCR cycles (Supplemental Tables S11 and S12). Thus, as long as the PCR product is sufficient for library pooling, we recommend to apply as few PCR cycles as possible and our PCR conditions for the current OSPALC protocol are: *<u>98 °C for 5 min followed by 10 cycles of 98 °C for 30 s, 55 °C for 30 s and 72 °C for 50 s, 10 cycles of 98 °C for 30 s, gradient 55°C to 65 °C for 30 s and 72 °C for 50 s, and one final extension step at 72 °C for 7 min</u>*.

### Applications of OSPALC to RNA samples from disturbed and stable environments

Because environmental DNA is highly stable, amplicon library constructions using DNA as templates could lead to false positives due to remnant DNA from dead organisms. RNA is transient and degrades quickly in the environment, and amplicon analyses based on rRNA could thus reveal the extant biodiversity of samples. Therefore, we applied OSPALC to total RNA extracted from environmental samples to investigate the dynamics of biodiversity in a pool before and after a hose-cleaning. Such application will test the performance of the OSPALC method on natural samples. The pool is located in front of the Ocean University of China (OUC) Library, Yushan campus (samples labelled as T1-4; Supplemental Fig. S2; the pool area is around 10 m^2^ and 1 m deep). The pool was hose-cleaned and refilled with water on June 8^th^, 2021. We collected samples from four spots as replicates. Each sample contains 500 ml water with sediment from the bottom of each spot, using zip bags on a golf ball retriever. Samples were collected mostly every two weeks from May 21^st^ 2021 to October 16^th^, 2021 (Supplemental Tables S13 and S14; Supplemental Fig. S2). Before being cleaned, the pool was murky and green, full of micro-organisms that were visible even to naked eyes. After hose-cleaning, the pool was clear and started to become green one week later (Supplemental Fig. S2). For each thoroughly-mixed sample, 50 ml was centrifuged at 4 °C and total RNA of the cell pellets was extracted immediately after collections and reverse-transcribed to cDNA, which was used as the template for the OSPALC PCR for the 16S-rRNA-V4 region.

The results are highly consistent with previous reports on bacterial communities in summer resting freshwater, as Proteobacteria and Cyanobacteria are the dominant bacteria (Zhang et al. 2018). Interestingly, probably because of the pool’s proximity to the sea (862 m to the nearest coast), some typical marine bacteria—*Pirellula, NS9_marine_group, Hyphomonas*—are also detected in these freshwater samples (Fig. 3). The sea breeze or activities of other living organisms may account for the detection of marine bacteria.

**Figure 3.**
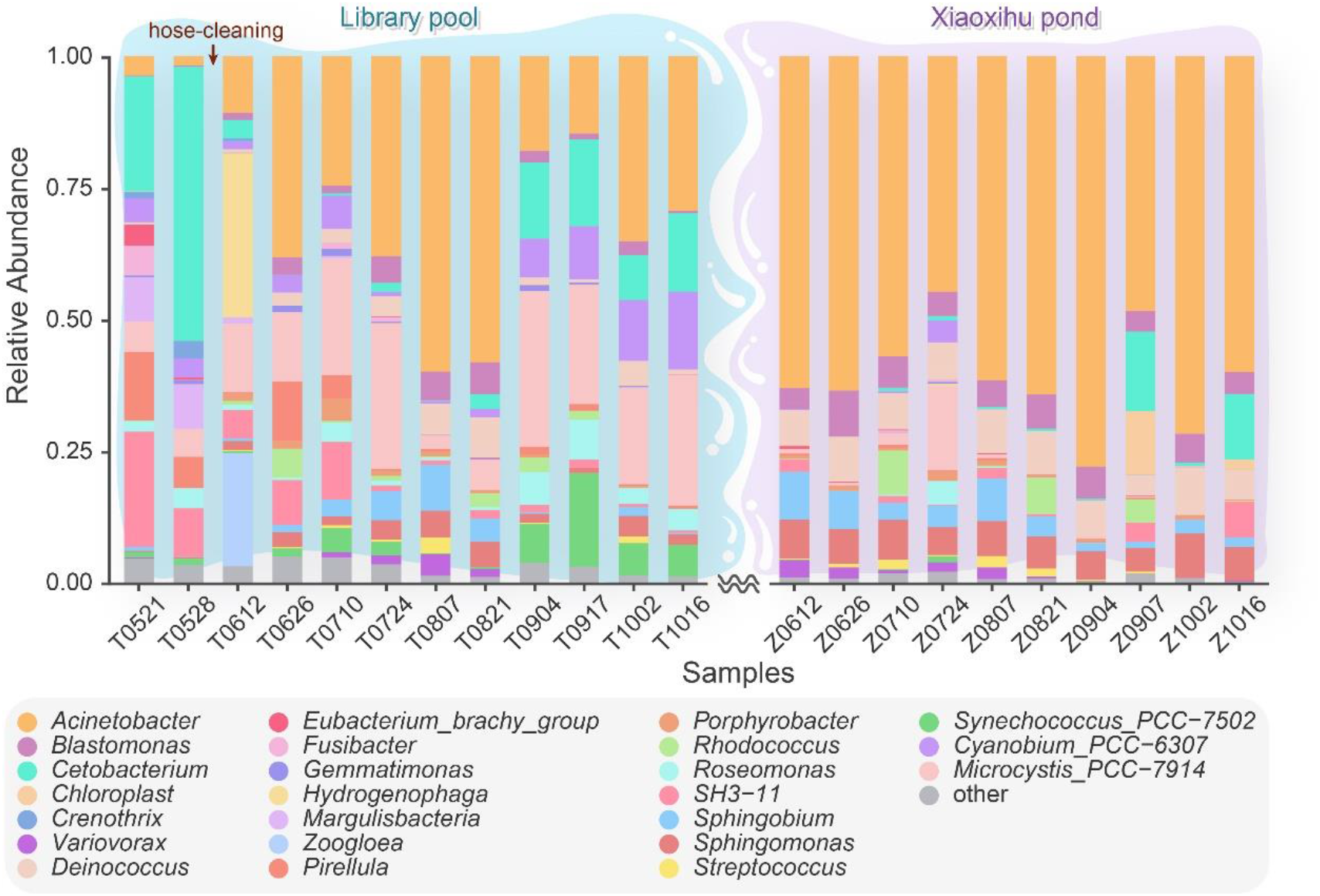
Relative abundance of the environmental samples from the two collection sites. The sequences of the four samples from each site on the same day were merged for analysis. T stands for the Library pool, Z stands for the Xiaoxihu pond in the Zhongshan park. The four digits after “T” or “Z” stand for the date in 2021. Library pool was hose-cleaned on June 8^th^, 2021.

The community composition, revealed by the amplicon sequencing, also shows clear-cut community successions, particularly for the samples collected before and after the hose-cleaning (June 8^th^, 2021; Fig 3; Supplemental Table S13). There was an explosive increase of the bacteria *Zoogloea* and *Hydrogenophaga* right after the environmental disturbance, but these almost disappeared in subsequent samples. *Zoogloea* are frequently reported in sewage or from sewage-treatment systems and *Hydrogenophaga* are able to perform dissimilatory nitrate reduction to ammonium (Dugan 1981; Tang et al. 2020). These bacteria may be opportunistic growers, particularly when competitors/predators are severely reduced after the hose-cleaning. Several species such as *Brevundimonas* and *Rhodococcus*, which were not detected before the hose-cleaning, also started to show up afterwards, with a low abundance. Several genera such as *Crenothrix* and *Margulisbacteria* gradually disappeared after the cleaning. *Crenothrix* are known to be important methane consumers (Oswald et al. 2017). These bacteria can be greatly affected by environmental changes such as the removal of leaves and sediments. From the phylum level, four days after the hose-cleaning, the proportion of Proteobacteria rose from 17.95% to 73.10%, and Fusobacteriota decreased from 43.30% to 3.18% (Fig. 3).

Besides the samples from an environment with perturbations, we also tested OSPALC on samples collected from another stable environment, Xiaoxihu pond, at Zhongshan Park of Qingdao, China (700 m to the above pool; area ~8000 m^2^, depth ~two meters, labelled as Z5-8). This pond has a history of 99 years and is rarely disturbed. We expected no extreme community successions around the hose-cleaning time of the above pool, and performed parallel sample collections with similar procedures as those for the above pool (Supplemental Tables S13 and S14). We then prepared OSPALC libraries using the total RNA extracted from the Xiaoxihu pond samples. To note, the PCR success rate of OSPALC could be elevated by diluting DNA to ~ 1ng/μl (data not shown), especially for the repeatedly-PCR-failed environmental samples. As expected, the community structure of the Xiaoxihu pond is stable across all sampling time points (Fig. 3). OSPALC could thus reliably reveal community structure and biodiversity results for environmental samples.

## Discussion

Compared with metagenomic sequencing, short-reads-based amplicon sequencing is not able to fully reveal the community function, since only part of the target gene is investigated (Myer et al. 2016). However, even the biodiversity indices and community compositions revealed from amplicon sequencing can be tremendously useful, compared with traditional methods using microscopy and staining (Eisenstein 2018; Gupta et al. 2019; Pochon et al. 2013). OSPALC can achieve this goal in a fast and cost-efficient way, compared with most amplicon library construction methods, especially those with the adapter ligation and indexing steps.

In principle, OSPALC could be applied to any conservative genes of most organisms, not limited to bacteria, contrastingly different from the service providers, which usually provide amplicon sequencing for 16S/18S rDNA only. The OSPALC results of the mock communities did not have contaminating species and generated fewer chimeras (Supplemental Tables S5– S7). As databases become more robust and more primer sequences are tested, amplicon sequencing is becoming more reliable and more widely-used in predictions based on Operational Taxonomic Units (OTUs), e.g. PICRUSt2 pathway prediction (Douglas et al. 2020). The extremely low cost, short hands-on time for just one regular PCR further makes OSPALC an appealing method. In addition, the full-length sequencing for 16S/18S rDNA amplicons using long-read sequencing is still expensive and error-prone compared with short-read sequencing, and in the same “genome fragment” status as partial gene amplicons do not confer an overwhelming advantage over short-read amplicons with greater resolution (Myer et al. 2016).

Nevertheless, with the continuous development of sequencing technology, full-length amplicons may become the mainstream method for studying community structure in the future. We have shown that PCR bias is not strongly associated with the deviation between genuine community composition and that from the amplicon analysis, and thus many other factors, such as GC content, copy number and so on can be the causes (Laursen et al. 2017). In order to solve these problems, we are developing a model that could hopefully calibrate the community composition deviation of OSPALC. The next version of OSPALC, which is being developed, will thus integrate cutting-edge sequencing technology, sophisticated algorithms, and target much larger genomic regions, from sequencing to data analysis.

## Methods

### Preparation of the two mock communities

Each mock community is a mixture of genomic DNA of five bacteria: two Gram-positive species—*Bacillus subtilis* ATCC6051, *Kocuria polaris* CGMCC1.8013; three Gram-negative species—*Escherichia coli* MG1655, *Pseudomonas aeruginosa* PAO1 and *Shewanella putrefaciens* CGMCC1.6515. All bacteria were cultured with conditions shown in Supplemental Table S4. The Marine LB Broth was prepared with a lab-developed recipe (Strauss et al. 2017): 1000 ml natural seawater with PSU of ~30, 5 g Bacto™ peptone (BD, Cat. No.: 9030688), 1 g Bacto™ yeast extract (BD, Cat. No.: 8344948), 0.18 g ferric-EDTA (SIGMA, Cat. No.: SLBV7746). DNA was extracted with the MasterPure™ Complete DNA and RNA Purification Kit (Epicentre, Cat. No.: MC85200). The Qubit 3.0 fluorometer and Nano-300 (Allsheng™) were used to measure the concentration and the purity of DNA, respectively. Four replicates were prepared for each mock community (Mock-Equal replicate 1 was discarded due to operation errors during mixing). *Kocuria polaris* CGMCC1.8013 genome was assembled by Unicycler 0.4.8 (Wick et al. 2017).

### Sample collection

Samples were collected from two sites in Qingdao, China: the pool in front of the library of Ocean University of China (Yushan Campus) and the Xiaoxihu pond at Zhongshan Park. About every two weeks (May 21^st^ – October 16^th^, 2021), we collected 500 ml bottom-water samples at each of the four spots of each site (Supplemental Fig. S2), using a golf-ball retriever with a zip bag. The OUC library pool experienced draining, hose-cleaning and refill (without using any disinfectant) on June 8^th^, 2021. The Xiaoxihu pond of Zhongshan Park was investigated as an un-disturbed comparison, and sampled similarly (Supplemental Fig. S2), which is much less humanly-disturbed as a site of view. After collection, each sample was immediately shipped to the lab, completely mixed and 50 ml sample was centrifuged at 4000 g and 4 °C. Temperature, dissolved oxygen, pH, salinity and conductivity were measured on site using one YSI Professional Plus Multiparameter Instrument (Supplemental Table S14).

### Nucleic acids extraction, amplicon library construction, and Illumina sequencing of environmental samples

After centrifugation of 50 ml of each sample, we discarded the supernatant and transferred the pellets in about 1.5 ml from the conical tube to 1.5 ml Eppendorf tubes. We then centrifuged again at 15000 g and 4 °C, for 3 min. RNA and DNA were extracted using the MasterPure™ Complete DNA and RNA Purification Kit (Cat. No.: MC85200). For reverse transcription of RNA to cDNA, we used Vazyme HiScript III 1st Strand cDNA Synthesis Kit (+gDNA wiper) (Cat. No.: R312). Details for all DNA or cDNA OSPALC library construction are in Supplemental File S1. Sterilized DI water was used as a control, and no contamination was detected after gel electrophoresis. Illumina Novaseq sequencing was performed at Berry Genomics, Beijing and Novogene, Tianjin.

### qPCR

We diluted genomic DNA of the five bacteria to 10 ng/μl, each with three replicates. With four serial 10-fold dilutions, five different concentrations of DNA for each bacterium were used for qPCR with Biometra GmbH 846-x-070-301 Biometra TOne 96G PCR machine. The 20 μl reaction system consisted of 2 μl DNA template, 10 μl AceQ^®^ Universal SYBR qPCR Master Mix (Cat. No.: Q511), 0.4 μl 515F primer (10 μM), 0.4 μl 806Y primer (10 μM) and 7.2 μl sterilized ultrapure water. The amplification conditions were: 95°C for 5 min followed by 40 cycles of 95 °C for 10 s and 60 °C for 25 s.

### Amplicon analysis

The amplicon analysis is based on QIIME2 (version: 2021.8) (Bolyen et al. 2019). Raw reads were joined with qiime vsearch (Rognes et al. 2016) and filtered by qiime q-score. The 16S rDNA V4 region database Silva was used—Silva 138 99% OTUs based on the 515F/806R sequences (Bokulich et al. 2018; Quast et al. 2012; Robeson et al. 2020). We performed feature classification with qiime deblur (Amir et al. 2017). We ran qiime alignment mafft for feature reads alignment and performed phylogenetic analysis with qiime phylogeny fasttree (Price et al. 2010). Overlapping bar charts were plotted with ggplot2 (Wickham 2016).

## Acknowledgments

This work was supported by the National Natural Science Foundation of China (31961123002, 31872228), the Fundamental Research Funds for the Central Universities of China (202041001), the Young Taishan Scholars Program of Shandong Province (tsqn201812024), National Science Foundation (DEB-1927159). We appreciate the technical help from Vazyme Biotech Co., Ltd., Nanjing and Wei Yang, OUC. All bioinformatic analyses were performed with IEMB-1 computation clusters at OUC. We also thank Haoyu Li, Jingjing Baoli and Zhirong Zhang for technical help.

