## Supplemental Figures S1-S2 for "One-Step PCR Amplicon Library Construction (OSPALC, version 1)"

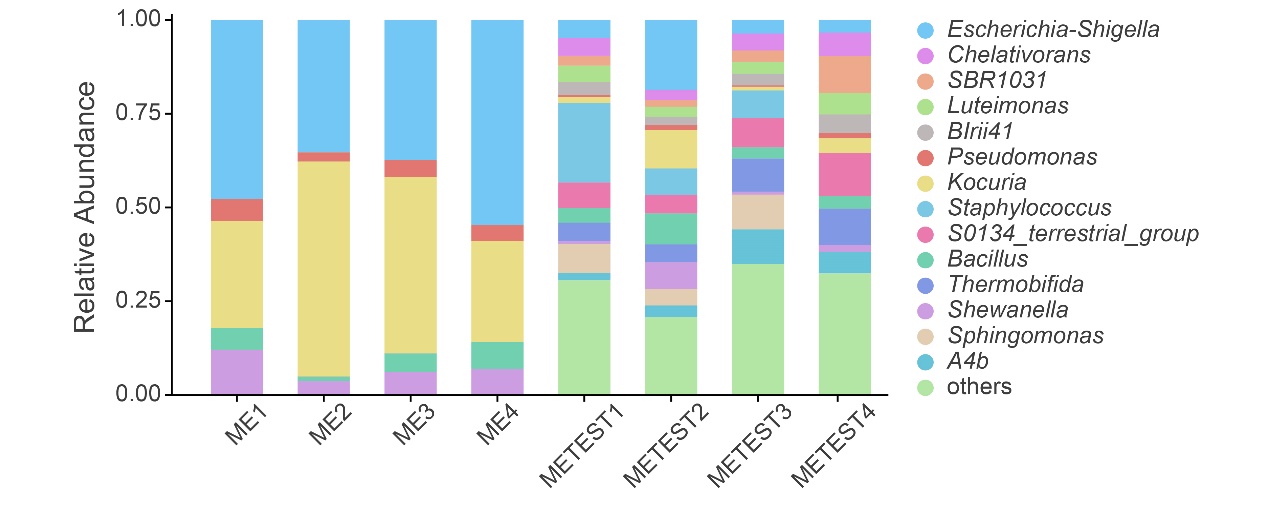

**Supplemental Figure S1.** Relative abundance of bacteria in the mock communities revealed by OSPALC (left four stacked bars; first batch of trials) vs. service provider (right four stacked bars; first batch of trials).

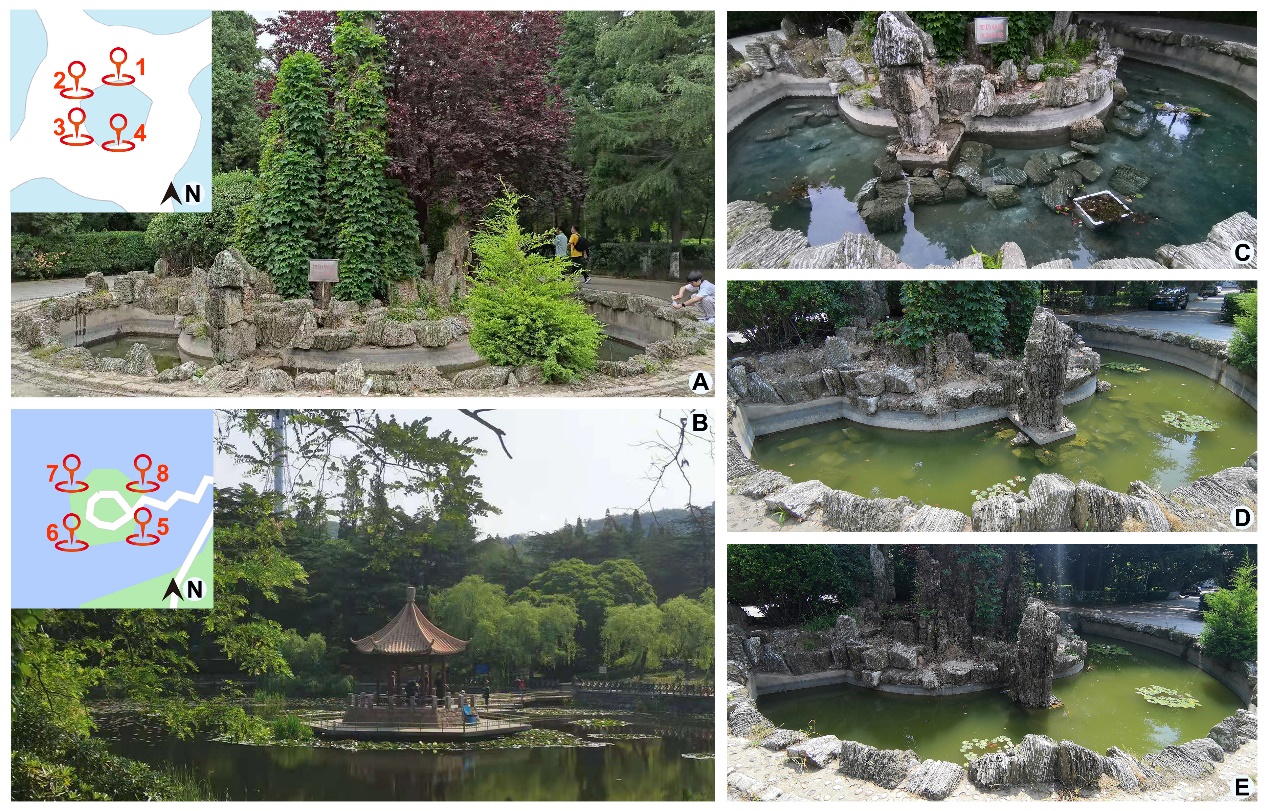

**Supplemental Figure S2.** **Collection sites and their sampling spots.** A: The OUC Library pool, the inset shows the four sampling spots in the pool; B: Xiaoxihu pond, the four sampling spots are in the inset; C, D, E: the OUC Library pool on June 8^th^, July10^th^ and August 7^th^, 2021.
