## Supplemental File S1 for "One-Step PCR Amplicon Library Construction (OSPALC, version 1)"

**Protocol for OSPALC library construction (10-μl reaction system)**

**2022-10-24**

**Main reagents**

- Vazyme 2× Phanta Flash Master Mix (Dye Plus) (Cat. No.: P520-02) (or any PCR high-fidelity kit)
- Primers 16SV4F1~16SV4F8
- Primers 16SV4R1~16SV4R12
- Sterilized ultrapure water
- OMEGA E.Z.N.A.^®^ Gel Extraction Kit (Cat. No.: D2500-02) (or any gel extraction kit)

**Equipment and Consumables**

- Jena Biometra GmbH 846-x-070-301 Biometra TOne 96G PCR machine (or any PCR machine)
- Multi-channel pipettes (2, 10, 200μl) + filter tips (always change tips!)
- 0.2ml 96-well PCR Plate (AXYGEN, Cat. No.: PMI0110-07A)
- Microseal 'B' PCR Plate Sealing Film, adhesive, optical (Bio-Rad, Cat. No.: MSB1001)
- NEST PCR sealing film (NEST, Cat. No.: 410001)
- Qubit^TM^ 3 Fluorometer (Invitrogen, Cat. No.: Q33216)
- Qubit^TM^ assay tubes (Invitrogen, Cat. No.: Q32856)

**library construction PROCEDURES**

**Prepare genomic DNA/environmental DNA for OSPALC library construction**

1. Dilute each DNA sample to 1ng/μl with nuclease-free water using a 96-well PCR plate; avoid cross-sample contamination; keep the PCR plate on ice. Sterilized DI water is used as control.

**PCR reaction system (10μl)**

2. Components of the reaction system for each sample:

Vazyme 2× Phanta Flash Master Mix (Dye Plus)	5.0μl

Forward primer	0.2μl

Reverse primer	0.2μl

Sterilized ultrapure water	2.6μl

DNA	2μl (1ng/μl)

**Total volume	10.0μl**

3. Prepare master-mix for 96 samples:

Prepare row master-mix in 8-strip PCR tubes A:

Forward primer 16SV4F1: 0.2×1.3×12=3.12μl, sterilized ultrapure water: 1.3×1.3×12=20.28μl, Vazyme 2× Phanta Flash Master Mix (Dye Plus): 2.5×1.3×12=39.0μl

Forward primer 16SV4F2: 0.2×1.3×12=3.12μl, sterilized ultrapure water: 1.3×1.3×12=20.28μl, Vazyme 2× Phanta Flash Master Mix (Dye Plus): 2.5×1.3×12=39.0μl

Forward primer 16SV4F3: 0.2×1.3×12=3.12μl, sterilized ultrapure water: 1.3×1.3×12=20.28μl, Vazyme 2× Phanta Flash Master Mix (Dye Plus): 2.5×1.3×12=39.0μl

Forward primer 16SV4F4: 0.2×1.3×12=3.12μl, sterilized ultrapure water: 1.3×1.3×12=20.28μl, Vazyme 2× Phanta Flash Master Mix (Dye Plus): 2.5×1.3×12=39.0μl

Forward primer 16SV4F5: 0.2×1.3×12=3.12μl, sterilized ultrapure water: 1.3×1.3×12=20.28μl, Vazyme 2× Phanta Flash Master Mix (Dye Plus): 2.5×1.3×12=39.0μl

Forward primer 16SV4F6: 0.2×1.3×12=3.12μl, sterilized ultrapure water: 1.3×1.3×12=20.28μl, Vazyme 2× Phanta Flash Master Mix (Dye Plus): 2.5×1.3×12=39.0μl

Forward primer 16SV4F7: 0.2×1.3×12=3.12μl, sterilized ultrapure water: 1.3×1.3×12=20.28μl, Vazyme 2× Phanta Flash Master Mix (Dye Plus): 2.5×1.3×12=39.0μl

Forward primer 16SV4F8: 0.2×1.3×12=3.12μl, sterilized ultrapure water: 1.3×1.3×12=20.28μl, Vazyme 2× Phanta Flash Master Mix (Dye Plus): 2.5×1.3×12=39.0μl

Prepare column master-mix in 12-strip PCR tubes B:

Reverse primer 16SV4R1: 0.2×1.3×8=2.08μl, sterilized ultrapure water: 1.3×1.3×8=13.52μl, Vazyme 2× Phanta Flash Master Mix (Dye Plus): 2.5×1.3×8=26.0μl

Reverse primer 16SV4R2: 0.2×1.3×8=2.08μl, sterilized ultrapure water: 1.3×1.3×8=13.52μl, Vazyme 2× Phanta Flash Master Mix (Dye Plus): 2.5×1.3×8=26.0μl

Reverse primer 16SV4R3: 0.2×1.3×8=2.08μl, sterilized ultrapure water: 1.3×1.3×8=13.52μl, Vazyme 2× Phanta Flash Master Mix (Dye Plus): 2.5×1.3×8=26.0μl

Reverse primer 16SV4R4: 0.2×1.3×8=2.08μl, sterilized ultrapure water: 1.3×1.3×8=13.52μl, Vazyme 2× Phanta Flash Master Mix (Dye Plus): 2.5×1.3×8=26.0μl

Reverse primer 16SV4R5: 0.2×1.3×8=2.08μl, sterilized ultrapure water: 1.3×1.3×8=13.52μl, Vazyme 2× Phanta Flash Master Mix (Dye Plus): 2.5×1.3×8=26.0μl

Reverse primer 16SV4R6: 0.2×1.3×8=2.08μl, sterilized ultrapure water: 1.3×1.3×8=13.52μl, Vazyme 2× Phanta Flash Master Mix (Dye Plus): 2.5×1.3×8=26.0μl

Reverse primer 16SV4R7: 0.2×1.3×8=2.08μl, sterilized ultrapure water: 1.3×1.3×8=13.52μl, Vazyme 2× Phanta Flash Master Mix (Dye Plus): 2.5×1.3×8=26.0μl

Reverse primer 16SV4R8: 0.2×1.3×8=2.08μl, sterilized ultrapure water: 1.3×1.3×8=13.52μl, Vazyme 2× Phanta Flash Master Mix (Dye Plus): 2.5×1.3×8=26.0μl

Reverse primer 16SV4R9: 0.2×1.3×8=2.08μl, sterilized ultrapure water: 1.3×1.3×8=13.52μl, Vazyme 2× Phanta Flash Master Mix (Dye Plus): 2.5×1.3×8=26.0μl

Reverse primer 16SV4R10: 0.2×1.3×8=2.08μl, sterilized ultrapure water: 1.3×1.3×8=13.52μl, Vazyme 2× Phanta Flash Master Mix (Dye Plus): 2.5×1.3×8=26.0μl

Reverse primer 16SV4R11: 0.2×1.3×8=2.08μl, sterilized ultrapure water: 1.3×1.3×8=13.52μl, Vazyme 2× Phanta Flash Master Mix (Dye Plus): 2.5×1.3×8=26.0μl

Reverse primer 16SV4R12: 0.2×1.3×8=2.08μl, sterilized ultrapure water: 1.3×1.3×8=13.52μl, Vazyme 2× Phanta Flash Master Mix (Dye Plus): 2.5×1.3×8=26.0μl

“1.3”is a magnification factor in case of reagents sticking to tips or pipette during transfers; keep the tubes on ice.

4. Prepare a new 96-well plate on ice, transfer 4µl row master-mix to wells in each row with a multichannel pipette from the 8-strip PCR tubes A, so that wells in the same row receive the same forward index.
5. Transfer 4µl column master-mix to wells in each column with a multichannel pipette from the 12-strip PCR tubes B, so that wells in the same column receive the same reverse index.
6. Transfer 2µl diluted DNA (1ng/μl) to each well.
7. Cover the 96-well PCR plate with a BioRad B sealing film tightly, especially press the edges to avoid leaking when heated in the PCR machine. Place the 96-well plate in the PCR machine and run the following program after preheat:

**Temperature	Time	Cycles**

105°C	Lid

98°C	Hold

98°C	5 min

98°C	30 sec

55°C	30 sec	10

72°C	50 sec

98°C	30 sec

55°C to 65°C	30 sec	10

72°C	50 sec

72°C	7 min

4°C	Hold

**Quality control and pooling of the libraries (i.e. PCR products)**

8. Take 1μl from each well of the 10µl 96-well plate and measure concentration by Qubit, assuming samples with concentration readings > 5ng/μl as successful ones.
9.
3. Purify the libraries for the target size (the size of the amplified gene fragments and both adaptors), using the OMEGA E.Z.N.A.^®^ Gel Extraction Kit (Cat. No.: D2500-02) or any other gel extraction kit.
10. Pool all the libraries together based on the concentrations measured by Qubit, with equal DNA amount for each sample and mean volume around 1μl.
11. Send ~20μl of pooled libraries, which are gel purified, to the service provider for PE250 sequencing. 100,000 reads (50,000 forward and 50,000 reverse reads) for each sample.
12. Cover the 96-well plate with the remaining libraries with a Nest PCR sealing film and store at -20°C (-80°C if for long-term storage).
